# New mitochondrial primers for metabarcoding of insects, designed and evaluated using *in silico* methods

**DOI:** 10.1101/325688

**Authors:** Daniel Marquina, Anders F. Andersson, Fredrik Ronquist

**Affiliations:** Department of Bioinformatics and Genetics, Swedish Museum of Natural History, SE-10405, Stockholm, Sweden.; Department of Zoology, Stockholm University, SE-10691 Stockholm, Sweden.; KTH Royal Institute of Technology, Science for Life Laboratory School of Biotechnology, SE-10691, Stockholm, Sweden.

**Author notes:** Corresponding author: Daniel Marquina; Department of Bioinformatics and Genetics, Swedish Museum of Natural History, SE-10405, Stockholm, Sweden;.

**Keywords:** *in silico* PCR, Hexapoda, primer bias, taxonomic resolution, DNA barcoding

## Abstract

Insect metabarcoding has been mainly based on PCR amplification of short fragments within the ‘barcoding region’ of the gene COI. However, because of the variability of this gene, it has been difficult to design good universal PCR primers. Most primers used today are associated with gaps in the taxonomic coverage or amplification biases that make the results less reliable and impede the detection of species that are present in the sample. We identify new primers for insect metabarcoding using computational approaches (ECOPRIMERS and DEGEPRIME) applied to the most comprehensive reference databases of mitochondrial genomes of Hexapoda assembled to date. New primers are evaluated in silico against previously published primers in terms of taxonomic coverage and resolution of the corresponding amplicons. For the latter criterion, we propose a new index, exclusive taxonomic resolution, which is a more biologically meaningful measure than the standard index used today. Our results show that the best markers are found in the ribosomal RNA genes (12S and 16S); they resolve about 90% of the genetically distinct species in the reference database. Some markers in protein-coding genes provide similar performance but only at much higher levels of primer degeneracy. Combining two of the best individual markers improves the effective taxonomic resolution with up to 10%. The resolution is strongly dependent on insect taxon: COI primers detect 40% of Hymenoptera, while 12S primers detect 12% of Collembola. Our results indicate that amplicon-based metabarcoding of insect samples can be improved by choosing other primers than those commonly used today.

## Introduction

Species identification based on sequencing of standard genetic markers — DNA barcoding — is now well established. The so-called ‘Folmer region’ of the cytochrome oxidase I (COI) gene of the mitochondrion (Folmer *et al.* 1994) has been widely accepted as the standard barcoding marker for Metazoa (Hebert *et al.* 2003), and we now have extensive reference libraries for many groups of organisms (CBOL database, http://www.boldsystems.org). A range of other markers are also used for DNA barcoding of animals. For instance, 16S (a mitochondrial ribosomal RNA (rRNA) gene) has been used for amphibians (Vences *et al.* 2005); 16S, 12S (another mitochondrial rRNA gene) or cytochrome b (CytB; a protein-coding mitochondorial gene) for fishes (Sevilla *et al.* 2007, Cawthorn *et al.* 2012) and more recently, an unexplored region at the 3’ end of the COI gene for odonates (dragonflies and damselflies) (Rach *et al.* 2017). However, the reference libraries for these alternative markers are small in comparison with that for COI.

In recent years, new High Throughput Sequencing (HTS) platforms have opened up the possibility of analyzing the taxonomic composition of entire environmental samples in a single analysis — metabarcoding. This has created a lot of excitement in the biodiversity research community. Unfortunately, the Folmer region of COI is not suitable for HTS platforms because it is too long. Therefore, special mini-barcodes (short fragments, of variable length and position, within the Folmer region) have been developed for metabarcoding. Hajibabaei *et al.* (2006) and Meusnier *et al.* (2008) showed that mini-barcodes from 135 bp up to ~ 450 bp can provide the same degree of taxonomic discrimination as the whole 658 bp Folmer region. Mini-barcodes have the additional advantage that they more easily can be amplified when the DNA is damaged or fragmented, which is common in environmental DNA samples (Taberlet *et al.* 2012, Yu *et al.* 2012).

Ideally, a marker used for metabarcoding should have highly conserved sequence stretches that can be used for the design of ‘universal’ primers amplifying all taxa of interest in the sample and that flank a highly variable region that can be used for species discrimination. Unfortunately, being a protein-coding gene, COI is highly variable in the third position of most codons due to the redundancy of the genetic code, making it quite challenging to design primers for metabarcoding with good taxonomic coverage (Deagle *et al.* 2014). Inevitably, there will be a varying number of mismatches between the primers and the templates in the sample, translating into differential affinity of the primers for different templates. The primer-template pairs with fewer mismatches will be amplified more easily in each cycle, potentially resulting in extreme overrepresentation of these sequences in the final PCR product. These biases in ‘universal’ COI primers have been documented empirically in several studies. Hajibabaei *et al.* (2011) and Brandon-Mong *et al.* (2015) reported biases with Lep-F1/Lep-R1 primers (Hebert *et al.* 2004), Yu *et al.* (2012) showed that the Folmer primers (Folmer *et al.* 1994) fail to amplify many species of Hymenoptera, and Clarke *et al.* (2014) showed that several primer pairs are associated with amplification bias resulting in overrepresentation of Diptera and Lepidoptera sequences. The use of degenerate primers can reduce the bias to some extent (Clarke *et al.* 2014, Elbrecht & Leese 2017). Morinière *et al.* (2016) studied the amplification success of four COI primer pairs with different degrees of degeneracy for different taxonomic groups taken from a Malaise trap sample. The amplification success was strongly dependent on degeneracy, varying from 5 *%* for primers with no degeneracy to 49 % for highly degenerate primers.

The amplification bias of COI primers has resulted in several metabarcoding studies exploring alternative markers. It is common, when working with a very broad taxonomic scope (up to phylum), to use a very conserved but easily amplified marker, such as the nuclear small subunit ribosomal RNA (rRNA) gene (18S) (Pawlowski *et al.* 2012). Several examples of 18S metabarcoding can be found, mostly involving soil/sediment biodiversity assessment (Drummond *et al.* 2015, soil; Brannock & Halanych 2015 and Dell’Anno *et al.* 2015, meiofauna) and eukoaryotic microorganisms (Pawlowski *et al.* 2012). For vertebrates, especially fishes, the most used alternative marker is the mitochondrial small subunit rRNA gene (12S), *e.g.* Valentini *et al.* (2016), Port *et al.* (2015) or Furlan *et al.* (2015). Performance of the mitochondrial large subunit rRNA gene (16S) has been tested in insect metabarcoding with promising results. Using *in silico* analyses, Clarke *et al.* (2014) showed that 16S mini-barcodes of less than 200 bp identified just slightly fewer species than mini-barcodes of COI of the same length when applied to a set of 315 species (constituting 264 genera and 23 orders) of insects, while the taxonomic coverage (no. of species successfully amplified) was 75-90 % with 16S versus only 50 % with COI. However, longer COI mini-barcodes increased the taxonomic resolution between closely related species to almost 100 %, while the resolution of 16S peaked at 85 %. Remarkably, taxonomic coverage and taxonomic resolution of 16S was consistent through 11 analyzed insect orders, while the best COI taxon coverage was just above 50 % within Diptera and Lepidoptera, and only between 0 % and 47 % within other insect orders.

Similar results were obtained by Elbrecht *et al.* (2016): 16S amplified more species and more equally through orders, thus enhancing biomass estimation. They state that if the goal is to identify the species present in the sample, COI is still the best choice due to the availability of extensive public reference databases, but when the aim is to assess the biodiversity in numbers — rather than in terms of species names — 16S would be a better choice. Additionally, 16S metabarcoding has the advantage of amplicons not being mistaken with nuclear pseudogenes or *Wolbachia*, as can be the case with COI (Clarke *et al.* 2014). Deagle *et al.* (2014) suggested that the best strategy to follow in the future would be to build local databases of several markers and conduct metabarcoding studies with these different markers simultaneously rather than focusing on a single universal marker (COI).

Clearly, there is a need for more systematic search for optimal metabarcoding markers. The increased interest in sequencing mitochondrial genomes in recent years, and the development of sophisticated software for primer design and evaluation, have opened up new and faster ways of tackling this task. Here, we take advantage of these opportunities in searching for optimal metabarcoding primers. Specifically, we compile all publicly available insect mitogenomes, and then apply two different primer design software packages, ECOPRIMERS and DEGEPRIME, to identify suitable markers for insect metabarcoding. ECOPRIMERS (Riaz *et al.* 2011; https://git.metabarcoding.org/obitools/ecoprimers/wikis/home) is part of the OBITOOLS bioinformatics package (Boyer *et al.* 2016; https://git.metabarcoding.org/obitools/obitools/wikis/home). Given amplicon length interval and maximum number of errors between primer and template (with the possibility of constraining those errors not to be in the 3’ end of the primer), ECOPRIMERS creates a list of the best primers from a set of sequences or genomes in the dedicated ECOPCR database format. DEGEPRIME (Hugerth *et al.* 2014a; https://github.com/EnvGen/DEGEPRIME) is a program for semi-automated design of degenerate primers, originally developed for microbial 16S rRNA metabarcoding but applicable more generally. Given a maximum degeneracy (number of unique primer sequence combinations) and primer length, DEGEPRIME creates a list of the best primers for an input alignment.

After identifying a set of suitable markers for insect metabarcoding using this approach, we then evaluate their performance against that of previously published ones, focusing on taxonomic coverage and resolution of species in the mitogenome reference database. We also assess the performance of different primer-pair combinations. Our results show that most previously published primers perform poorly compared with the optimal primers or primer-pair combinations identified here.

## Materials and Methods

### Data preparation

Two sets of complete mitochondrial genomes of Hexapoda were created by downloading all accessible entries from GenBank, the first accessed in October 2015 and the second accessed in September 2016 (Fig. S1; Tables S1 and S2). The first set (D1), comprising a total of 1138 genomes (corresponding to 801 species, 607 genera, 268 families and 34 orders; 75 species (9 %) were represented by more than one sequence), was converted into ECOPCR database format (for use with ECOPRIMERS) and, in parallel, split into different fasta files, each containing either a protein coding gene or an rRNA gene, using GENEIOUS 8.1.7 (https://www.geneious.com, Kearse *et al.* 2012) (for use with DEGEPRIME). The protein-coding genes ATP8, ND2 and ND6 could not be extracted by aligning them to a reference sequence using the default settings of GENEIOUS due to high variability. Consequently, they were not considered for designing primers that could amplify a wide range of insect taxa. The rest of the extracted genes (12S, 16S, ATP6, COI, COII, COIII, ND1, ND3, ND4, ND4L and ND5) were aligned using MAFFT v7.266 (Katoh & Standley 2013). The second set (D2), comprising a total of 1600 genomes (corresponding to 1081 species, 766 genera, 311 families and 34 orders; 948 of the species (12 %) were represented by more than one sequence) was converted into ECOPCR database format. D1 was used for primer design and D2 for primer evaluation. To assess the accuracy of the taxonomic annotation of the GenBank entries, the complete COI sequence of each entry in D1 was submitted to BOLD database using custom scripts. The GenBank species identification for each sequence and the identity provided by BOLD were then compared.

### Design of new primers

Primer design was done using dataset D1. The script TrimAlignment.pl (included in the DEGEPRIME package) was run over the sets of extracted genes, trimming away alignment columns with more than 10 % of the sequences having gaps (`-min 0.9`).

For the DEGEPRIME analysis, we used a primer length of 18 bp (`-l 18`) and maximum degeneracy of 12- 216-fold (`-d 12 / -d 216`). Maximum degeneracy was set low (12-fold) to find primers with high specificity and low risk of forming primer dimers, and higher (216-fold) to explore results from the other end of the trade-off between unspecificity/primer dimers and higher sequence matching. These analyses will be referred to as DEGEPRIME-d12 (12-fold degeneracy) and DEGEPRIME-d216 (216-fold degeneracy). Entropy for each potential primer site is calculated by DEGEPRIME as ∑*p*_*i*_ log_2_ *p*_*i*_, where *p*_*i*_ is the frequency of sequence *i*, where sequence *i* has the same length as the primer. After finding primer sites with low entropy, we identified the best primer pairs for each mitochondrial gene amplifying a sequence of suitable length for metabarcoding (100-500 bp long). ECOPRIMERS was run over the ECOPCR database formatted dataset with the options of amplicon length of 50-500 bp (`-l 50 -L 500`), no mismatches in at least 70 % of the species (default) and up to three mismatches in 90 % of the species (default), none of them in the 3’ end of the primer (`-3 3`) and considering the sequences as circular (since the mitochondrial genome is circular) (`-c`).

### In silico *PCR*

The primers found in the previous step, plus already published primers for barcoding and metabarcoding of insects targeting COI, 16S and CytB, were subjected to in silico PCR using D2 and ECOPCR in the OBITOOLS package (Ficetola *et al.* 2010; https://git.metabarcoding.org/obitools/ecopcr/wikis/home). Compared to D1, D2 contains approximately 500 more sequences and 200 more species, thus providing a good test of the ability of the primers to detect and discriminate new taxa. Stringent PCR conditions (corresponding to high annealing temperature, resulting in higher specificity and lower amplification bias) were emulated by setting the ECOPCR options such that we allowed no mismatches (`-e 0`), and amplicon length ± 10 % of the expected length for each primer pair. Nine insect orders within D2 were also amplified separately. Specifically, we targeted those insect orders that are most abundant in Malaise traps (Hymenoptera, Diptera, Lepidoptera, Hemiptera) and in pitfall traps (Coleoptera, Collembola, Blattodea, Orthoptera and Thysanoptera).

### Measuring the performance of markers

The performance of each marker, consisting of a primer pair and its associated amplicon, was assessed using different indices measuring taxonomic coverage and resolution. *Taxonomic coverage* (*B*_*C*_) is the proportion of species in the dataset that are amplified by the given primer set (Ficetola *et al.* 2010). *Taxonomic resolution* (*B*_*S*_) is the proportion of species whose amplified sequences are unambiguously separated from the remainder of the set at a given similarity threshold (Ficetola *et al.* 2010). A species is considered unambiguously identified when it does not present any synonymy conflict, that is, when no cluster of barcodes from the species contains sequences of other species. However, this way of measuring resolution does not penalize the presence of two or more clusters with the same species label. This is not a problem when the analysis is reference-based, *i.e.* the downstream diversity analysis aggregates split clusters based on taxonomic annotations in a reference database. However, when such a reference database is missing and the analysis is focused on molecular operational taxonomic units (MOTUs), this measure artificially inflates taxonomic resolution as the similarity threshold increases (Riaz 2011). To address MOTU-based scenarios, we propose an alternative measure of taxonomic resolution, which we refer to as *exclusive taxonomic resolution* (*B*_*E*_), in which the presence of two or more clusters sharing the same species label is considered as an ambiguity (even when each of these clusters only contains sequences with the same species label).

It is important to note that *B*_*S*_ and *B*_*E*_ are calculated over the set of sequences amplified by the primer pair, not over the original set, which can potentially lead to misinterpretations in case of high values of *B*_*S*_ (or *B*_*E*_) and low values of *B*_*C*_. For instance, if a primer pair amplifies 60 % of the species in a mixture and the generated barcodes are able to discriminate between 87 % of those amplified species, one might get the impression of having a good primer pair, while the reality is that such a primer pair is incapable of detecting almost half the species in the sample (87 % of 60 % is 52 %). To get a general idea of how many species can be detected in a sample, we propose the *effective taxonomic resolution*(*ETR*) index, defined as the product between *B*_*C*_ and *B*_*E*_.

For a formal definition of these measures, let *S*_*U*_ be the set of species occurring in a single, homogeneously labeled cluster (a uniquely resolved or unambiguously identified species) and let *S*_*R*_ be the set of species occurring in a homogeneously labeled cluster (regardless of whether there are more clusters with the same label). Let *S*_*A*_ be the set of species among the amplified sequences and let *S* be the total set of species in the database. With standard set theory notation, where |*S*| denotes the number of (unique) elements in a set *S*, we can then define the indices as follows: *B*_*C*_ = |*S*_*A*_| / |*S*|, *B*_*S*_ = |*S*_*R*_| / |*S*_*A*_|, *B*_*E*_ = |*S*_*U*_| / |*S*_*A*_|, and *ETR* = *B*_*C*_*B*_*E*_ = |*S*_*U*_**|**/|*S*|.

An important property of *B*_*E*_ is that it varies with the similarity threshold used for clustering, peaking at a value (the ‘barcoding gap’) that is characteristic for the marker. If the similarity threshold is too low, many closely related species will not be distinguished; if it is too high, variable species will lower *B*_*E*_. To identify this peak in *B*_*E*_, it is important to have many species represented by multiple sequences. Our reference databases have many singletons, and only a few species represented by multiple sequences. To facilitate the identification of the optimal similarity threshold under such circumstances, we introduce an alternative definition of exclusive taxonomic resolution, 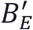, which is defined relative to clusters and not species. Thus, it penalizes oversplitting by counting the additional clusters generated by splitting the species that are well represented in the database into more than two clusters, using this to compensate for the fact that we cannot detect oversplitting in the many singleton species.

At peak resolution (the barcoding gap), 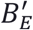 should be a reasonable approximation of *B*_*E*_ measured over a database where all species are represented by many sequences. For a formal definition, let *C*_*A*_ be the total set of clusters produced during the clustering of the amplified sequences. This set is composed of three subsets: *C*_*U*_ is the set of clusters with a single, unique species label, *C*_*M*_ is the set of clusters with more than one label and *C*_*N*_ is the set of clusters with a single but not unique label (*i.e.*, the label is shared with other clusters). Then, we define the alternative index 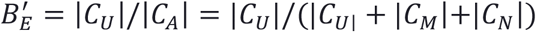. Note that |*C*_*U*_| = |*S*_*U*_| but |*C*_*A*_| ≥ **|***S*_*A*_**|**.

### Combinations of primer pairs

Two approaches were used to find good combinations of markers (primer pairs and associated amplicons) for metabarcoding. Firstly, we simply examined pairs of markers that were identified in the previous steps as having good performance when used on their own (independent approach). Secondly, we considered the best marker identified for each gene in the previous steps, and then searched for the best marker for the fraction of the dataset that the first marker was unable to detect (residual approach).

To measure the success of a pair of markers, we looked at the total number of species resolved by at least one of the two markers relative to the total number of species in the database. We regarded this as the total effective taxonomic resolution of the two markers, *ETR*_*T*_. The contribution of each marker was then be teased apart by focusing on the species that were uniquely resolved by one marker but not the other. Formally, let 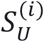 be the set of species uniquely resolved by marker *i*, with similar index notation for other species sets. Then we define the total effective taxonomic resolution of two markers *i* and *j* as

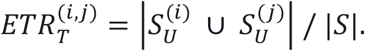

and the uniquely contributed taxonomic resolution of marker *i* as

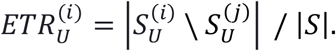

Note that *S*_*X*_\*S*_*Y*_ refers to the elements occurring in *S*_*X*_ but not in *S*_*Y*_. Finally, we define the redundant taxonomic resolution of two markers *i* and *j*, that is, the species that are unambiguously resolved by both markers, as

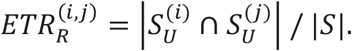

Note that these indices are additive, such that

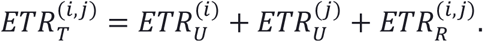

Primer quality indices and other definitions are summarized in Table 1. Unique, redundant and combined sets of species were computed using custom scripts.

**Table 1.** Primer and barcode quality indexes.

| Index | Symbol | Summary |
| --- | --- | --- |
| **Taxonomic coverage** | $B_C$ | Proportion of species whose sequences are amplified by the primer pair. |
| **Taxonomic resolution** | $B_S$ | Proportion of species whose amplified sequences are unambiguously identified, not considering repeated species labels as ambiguity. |
| **Exclusive taxonomic resolution** | $B_E$ | Proportion of species whose amplified sequences are unambiguously identified, considering repeated species labels as ambiguity.<sup>(1)</sup> |
| **Effective taxonomic resolution** | ETR | Proportion of species whose sequences are amplified and unambiguously identified, considering repeated species labels as ambiguity. |
| **Redundant effective taxonomic resolution** | $ETR_R$ | Effective taxonomic resolution shared by two or more barcoding markers applied to the same sample. |
| **Uniquely contributed effective taxonomic resolution** | $ETR_U$ | Effective taxonomic resolution obtained by just one of two or more simultaneous barcoding markers, but not by the other(s). |
| **Marker $j$ residual over marker $i$** | $jri / ETR_{jri}$ | Marker amplified by primers designed over the set of species not amplified by marker $i$. ETR of marker $jri$. |

<sup>(1)</sup> In this article, an alternative of $B_E$ is used due to the characteristics of the datasets, calculated in base of the number of clusters instead of species ( $B'_E$ )

### Computations

The *B*_*C*_ index values were computed using the command `ecotaxstat` from the OBITOOLS package. The *B*_*S*_ values were calculated using the command `ecotaxspecificity` with a similarity threshold varying from 95 % to 100 % in steps of 1 %. The same similarity threshold range was used with the algorithm UCLUST (implemented in the program USEARCH (Edgar 2010)), and then custom scripts were used to count the number of clusters with a single, unique species label, |*C*_*U*_|, and the clusters with mixed labels or with a single but not unique label, |*C*_*M*_| and |*C*_*N*_|, which were then used to compute 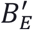 and all variants of the *ETR* index. All scripts used for the study are available at https://github.com/metagusano.

## Results

### Primer design

Potential primer sites in the rRNA genes (12S and 16S) have significantly lower entropy than the best primer sites in the protein-coding genes (Figs. 1, S2). Both rRNA genes offer primer sites with entropy well below 4, which is rare in the other genes. Only primer pairs matching more than half of the D1 sequences were considered for further examination. For DEGEPRIME-d12, no primers filling this requirement were found for ATP6, ND1, ND3, ND4 or ND4L, but a large number of potential primer pairs were identified in the remaining genes (Table S3). The 12S primers F1 and F2, as well as R1 and R2, are very closely located one to each other and partially overlap with SR-J-14199 - SR-N-14594 published by Kambhampati & Smith (1994). One of the 16S reverse primers (Hex16SR2) coincided with the reverse complement of the primer Ins16S_9R published by Clarke *et al.* (2014). For Cytochrome B, the forward primer of one of the pairs we found coincided with the reverse complement of REVCB2H of Simmons & Weller (2001), while the forward primer of the other pair corresponds to the same primer but shifted one position forward in relation to REVCB2H. None of the other primers we found correspond to previously published primers.

**Fig. 1.**
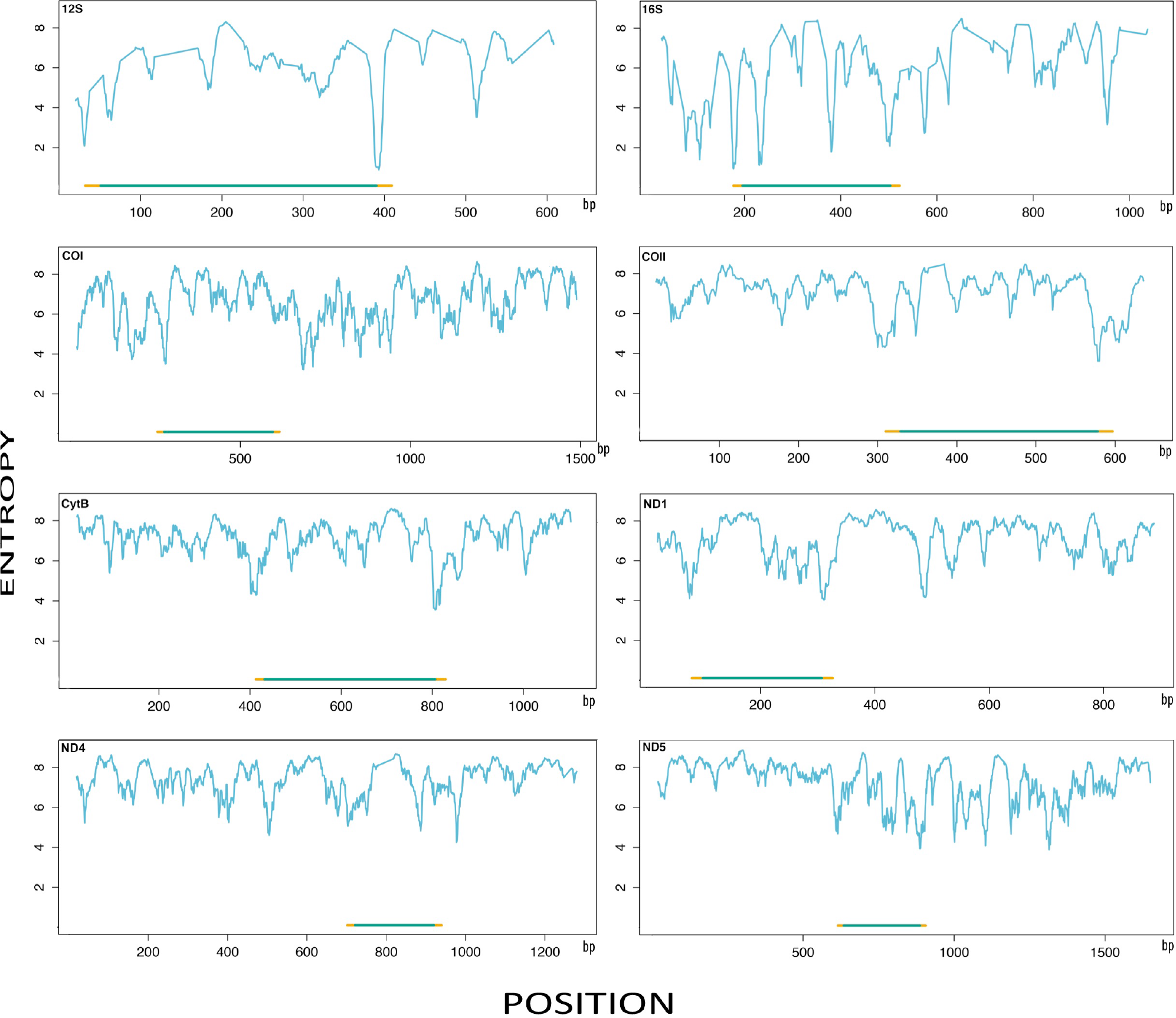
Entropy as a function of potential primer site (of length 18bp) in the eight mitochondrial genes that are best for metabarcoding. At the bottom of each figure, we give the position of the marker we selected for more detailed study. For each marker, the yellow lines represent the primer pair, and the green line the resulting amplicon.

For DEGEPRIME-d216 (Table S4), we found primers matching more than half of the D1 sequences in all genes except ND4L. A high proportion of DEGEPRIME-d216 primers are more degenerate versions of primers from DEGEPRIME-d12. All DEGEPRIME-d12 primers that partially overlapped with previously published primers (see above) had more degenerate DEGEPRIME-d216 versions. The COI primer pair HexCOIF4 - HexCOIR4 found in DEGEPRIME-d216 partially overlaps with BF2 – BR1 published by Elbrecth & Leese (2017) and ArF(1-5,10)-ArR(2,3,5,6,7,9) published by Gibson *et al.* 2014. Finally, the DEGEPRIME-d216 primer HexCytBF3 partially overlaps with the reverse complement of REVCB2H of Simmons & Weller (2001).

ECOPRIMERS found five primer pairs, which are all combinations between two forward primers and three reverse primers (F1-R1, F1-R2, F2-R2, F1-R3, F2-R1; see Table S5). All five pairs amplify fragments of the 16S gene.

### Measuring the quality of primer pairs

Among all primer pairs analyzed (found using DEGEPRIME, ECOPRIMERS or previously published), the ones with the highest coverage (*B*_*C*_) for each gene were used to test the differences between different ways of measuring resolution (*B*_*S*_ and 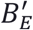). For all markers, taxonomic resolution measured in the standard way (*B*_*S*_) increased monotonically as the similarity threshold increased, reaching its maximum at 100 % (Fig. 2A), that is, when only identical sequences are considered to belong to the same cluster. The resolution was very similar for the different markers at 100 % similarity threshold (0.95-0.98), while differences increased at lower thresholds, ranging from 0.57 for 12S to 0.84 for COIII and CytB at a similarity of 95 %.

**Fig. 2.**
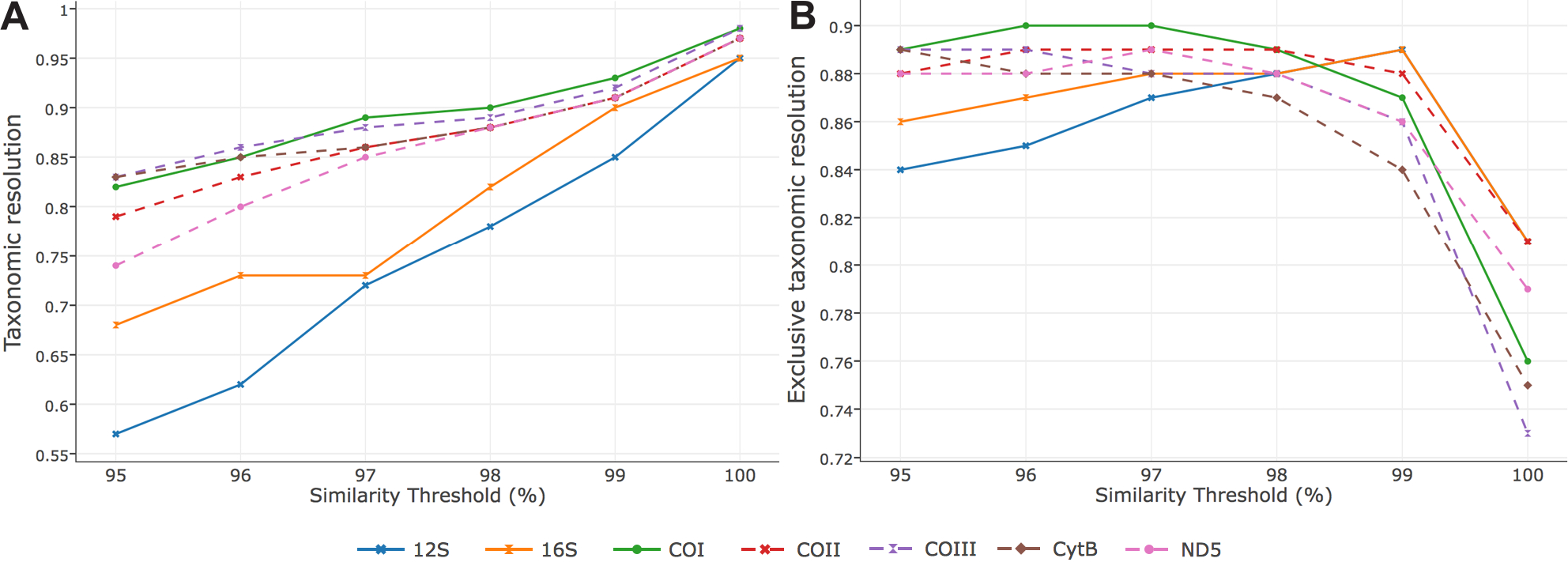
Taxonomic resolution (*B*_*S*_) (**A**) and exclusive taxonomic resolution (*B*_*E*_′) (**B**) as a function of the similarity threshold used in clustering sequences. For all genes, taxonomic resolution measured in the standard way increases as the similarity threshold increases, while exclusive taxonomic resolution peaks at intermediate levels of similarity, indicating where the barcoding gap is for each marker.

In contrast, exclusive taxonomic resolution 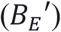 of different markers peaked at different intermediate similarity thresholds (Fig. 2B), the peak corresponding to where the barcoding gap stands for each one of them. Maximum 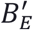 was reached at a similarity threshold of 99 % for 12S and 16S; 96-97 % for COI, 96-98 % for COII, 95-96 % for COIII, 95 % for CytB and 97 % for ND5. Measuring exclusive taxonomic resolution relative to species rather than clusters resulted in curves that were intermediate between *B*_*S*_ and 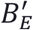, as expected (data not shown). These curves did not allow safe identification of the optimal similarity threshold values for different markers. In all subsequent analyses, the optimal similarity threshold value identified using 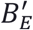 was chosen for each primer pair evaluated (97 % in the case of COI to comply with the common practice in metabarcoding, 97 % in the case of COII and 96 % in the case of COIII).

Note that the peak of exclusive taxonomic resolution is at 89-90 % of the species in the D2 dataset for all primers. Most of the unresolved 10 % of species are shared between different markers (Table S10), suggesting that only 89-90 % of species in D2 can be resolved using mitochondrial markers. The remaining species either have more intraspecific or less interspecific variation in their mitochondria than is typical, or the taxonomic annotation is incorrect. Given that the disagreement in the taxonomic annotation of D1 and D2 between GenBank and BOLD (considered to be better taxonomically curated) is around 10 % (Table S11), it seems likely that the main explanation is erroneous taxonomic labeling of the sequences.

### Performance of single markers

Figure 3 shows the performance of previously published markers (primer pair and associated amplicon) and the best markers designed with DEGEPRIME-d12, DEGEPRIME-d216 and ECOPRIMERS. Detailed results for all primers are given in the supplementary material (Tables S3-S5 and S8-S9; see also separate csv files S12-S16).

**Fig. 3.**
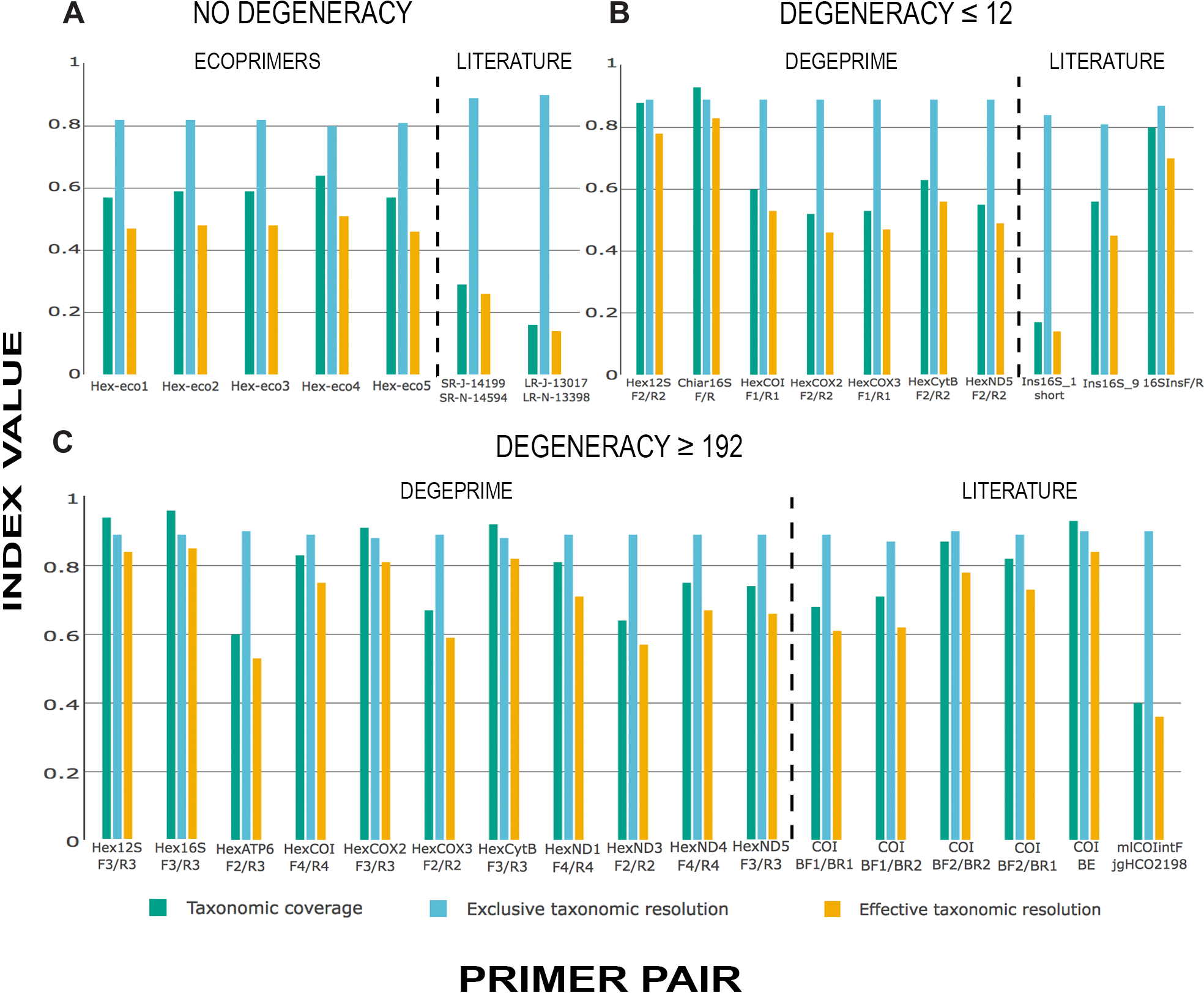
Taxonomic coverage (*B*_*C*_, green), exclusive taxonomic resolution (*B*_*E*_′ blue) and effective taxonomic resolution (*ETR*, orange) of examined primers. **A**. Newly designed primers with ECOPRIMERS and published primers with no degeneracy. **B**. Newly designed primers with DEGEPRIME and published primers with degeneracy lower or equal to 12-fold. **C**. Newly designed primers with DEGEPRIME and published primers with degeneracy higher than 192-fold. Among the published primers, only those with *B*_*C*_ > 0.15 are shown in this graph.

The DEGEPRIME-d12 markers (Fig. 3, Table S3) fall into two groups with respect to taxonomic coverage (*B*_*C*_): the two rRNA genes with high coverage (around 0.80-0.90), and the protein coding genes with intermediate levels of coverage (0.50–0.60). The taxonomic coverage roughly reflects the entropy of the primer pairs, low entropy corresponding to high taxonomic coverage. The taxonomic resolution 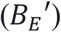, however, is more similar among markers, ranging from 0.80 for the worst 16S amplicon (Hex16SF2 - Hex16SR2) to 0.89 in the best markers (note that 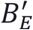 is calculated at different similarity thresholds depending on the variability of the gene). Thus, the markers with the broadest coverage are only slightly worse in distinguishing between sequences of closely related species, even though they need to consider a larger set of species. This results in the effective resolution being considerably higher for the rRNA markers (*ETR* = 0.71-0.83) than for the markers in the protein-coding genes (*ETR* = 0.46-0.53).

The more degenerate DEGEPRIME-d216 primers (Fig. 3, Table S4) for the 12S rRNA gene improved coverage considerably (from 0.81-0.88 to 0.94) compared to the DEGEPRIME-d12 primers, but there was only a slight increase for the 16S rRNA gene (from 0.92-0.93 to 0.96). For the protein-coding genes, the improvement was more striking. Relaxing the stringency to 216-fold degeneracy produced competitive primers for genes where adequate primers with 12-fold degeneracy did not exist (ATP6, ND1, ND3 and ND4). It also significantly increased the coverage of COI, COII and CytB primers, while the effect was smaller for COIII and ND5. The case of COII and CytB is especially noteworthy, since the degenerate primers for these genes reach *B*_*C*_ values as high as those of the best rRNA primers with 12-fold degeneracy (Fig. 3, Tables S3-S4).

The metabarcoding markers found using ECOPRIMERS (Fig. 3, Table S5) exclusively target regions of the 16S gene. However, they have considerably lower coverage (*B*_*C*_ = 0.57-0.64) and effective resolution (*ETR* = 0.46-0.51) than the 16S markers found with DEGEPRIME.

Among the already published primer pairs (Table S8), resolution of the corresponding amplicon is generally high 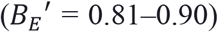, but the coverage varies over a wide range depending on the degeneracy of the primers. The 16S primer pairs that only include degenerate bases in the forward primer (Ins16S_1 and Ins16S_1short), or a single degenerate base in the reverse primer (Ins16S_9) have low to intermediate coverage values (*B*_*C*_ = 0.06-0.56), while the pair that has degeneracy in both primers (16SIns_F/Ins_R) has considerably higher coverage (*B*_*C*_ = 0.80). Effective taxonomic resolution (*ETR*) ranges from 0.14 (Ins16S_1short) to 0.70 (16SIns_F/Ins_R). For COI, the effect of primer degeneracy is even more pronounced, with moderate to high coverage (*B*_*C*_ = 0.68-0.87) for the highly degenerate primers BF/BR, while the coverage was much lower (*B*_*C*_ = 0.00-0.06) under the strict PCR settings used here for primers with low (only one degenerate base, or several degenerate bases but only in one primer of the pair) or no degeneracy (Table S8). As a consequence, *ETR* is medium to high (0.61-0.78) for the pairs involving BF/BR combinations, and close to zero for the remaining pairs. Coverage of the BE fragment of COI, amplified by the primers ArF2 - ArR5 (Gibson *et al.* 2014), is as high as with the rRNA genes (*B*_*C*_ = 0.93) when inosine is set to pair with all nucleotides (acting as an N), but was considerably reduced (*B*_*C*_ = 0.72) when set to pair with only A, T and C (acting as an H). The single primer pair targeting CytB (REVCB2H - REVCBJ), with only the forward primer with degeneracy of 2-fold, also had low taxonomic coverage (*B*_*C*_ = 0.01), and ETR close to zero.

Further evaluation was only performed for the best primer pair for each gene, the pair with the highest *ETR* value (Table 2). For all genes except COI, the best pair found was one designed using DEGEPRIME. For COI, the best pair found using DEGEPRIME was very similar to the pair BF2 - BR1 (ELbrecht & Leese 2017): both of the HexCOIF4 - HexCOIR4 primers are two bases shorter than the BF2 - BR1 primers, and there are two substitutions (Y for T in the 18^th^ base starting from the 5’ end and N for D in the 3^rd^ base starting from the 5’ end) that provide HexCOIF4 - HexCOIR4 with a higher coverage than BF2 - BR1 (0.75 versus 0.72). Although the combination BF2 - BR2 has even higher *ETR*, the amplicon length of BF2 - BR1 or HexCOIF4 - HexCOIR4 is more suitable for today’s sequencing platforms. In the end, we therefore selected HexCOIF4 - HexCOIR4 for further study. Also, given that the increase in *ETR* values between rRNA primers with 12-fold and 216-fold degeneracy is modest, and that the degenerate rRNA primers could potentially amplify non-insect DNA from environmental samples during the PCR because of the low variability of the rRNA gene, we selected the rRNA primers with 12-fold degeneracy for further study.

**Table 2.**
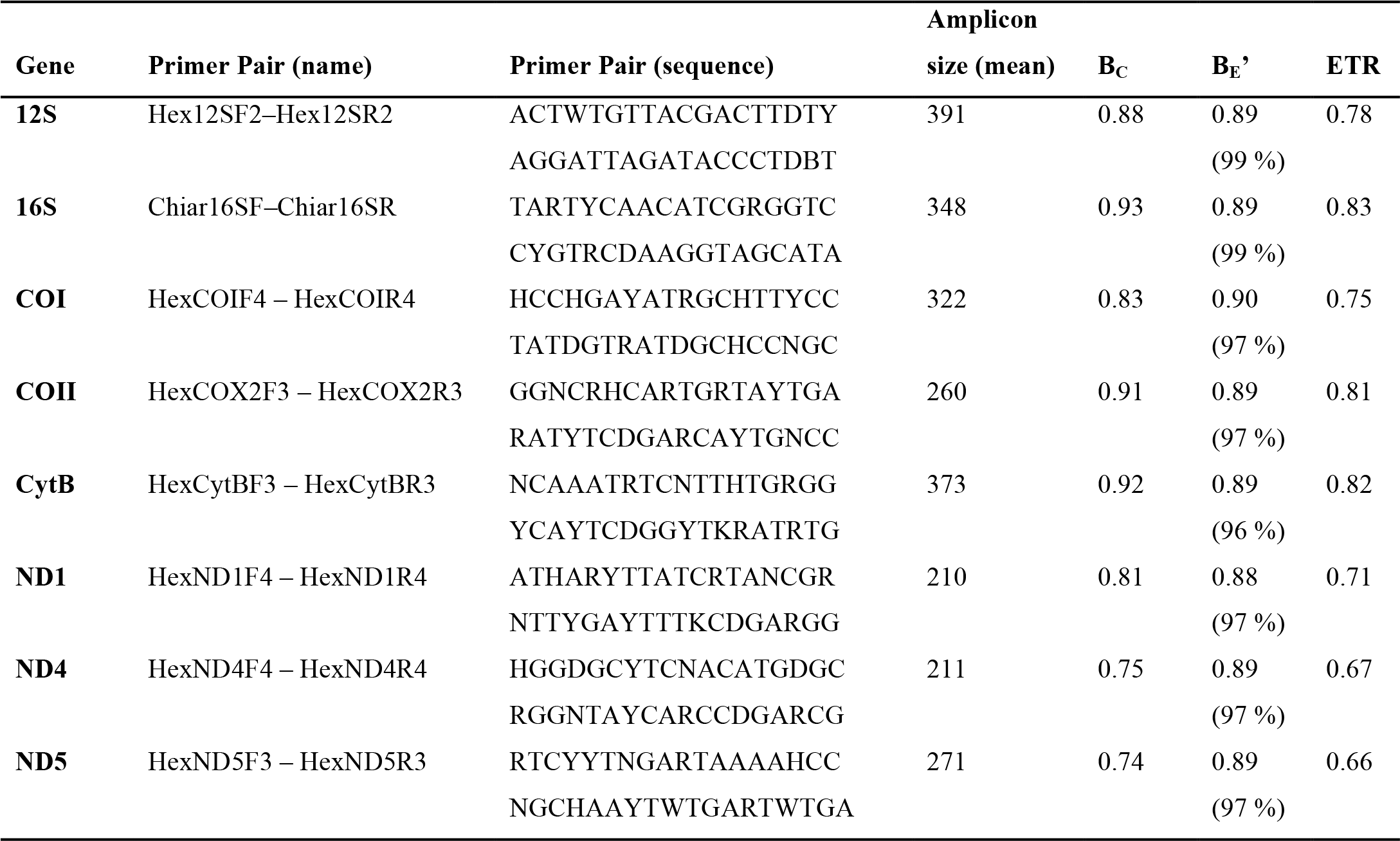
The best primers for insect metabarcoding found for each mitochondrial gene. Together with *B*_*E*_′, we give the optimal similarity threshold in parentheses for each marker.

Among the nine insect orders most abundantly found in Malaise traps and pitfall traps, the highest *ETR* is provided by 16S for Diptera, Coleoptera, and Collembola (Fig. 4). The highest *ETR* for Orthoptera and Hemiptera is provided by 12S, while CytB is the best marker for Coleoptera and Thysanoptera, and COII for Hymenoptera, Lepidoptera and Blattodea (ND5 provides equal *ETR* for Blattodea). Surprisingly, COI doesn’t provide the highest ETR for any of the nine orders, although it performs reasonably well for all of them except Hymenoptera and Thysanoptera. The only order for which 16S fails is Thysanoptera, while 12S provides low *ETR* for Thysanoptera, Collembola and Hymenoptera. Values of *ETR* of CytB and COII are also generally high with exceptions: COII fails for Thysanoptera and CytB provides low *ETR* for Collembola. The rest of the genes vary dramatically in performance across the nine orders.

**Fig. 4.**
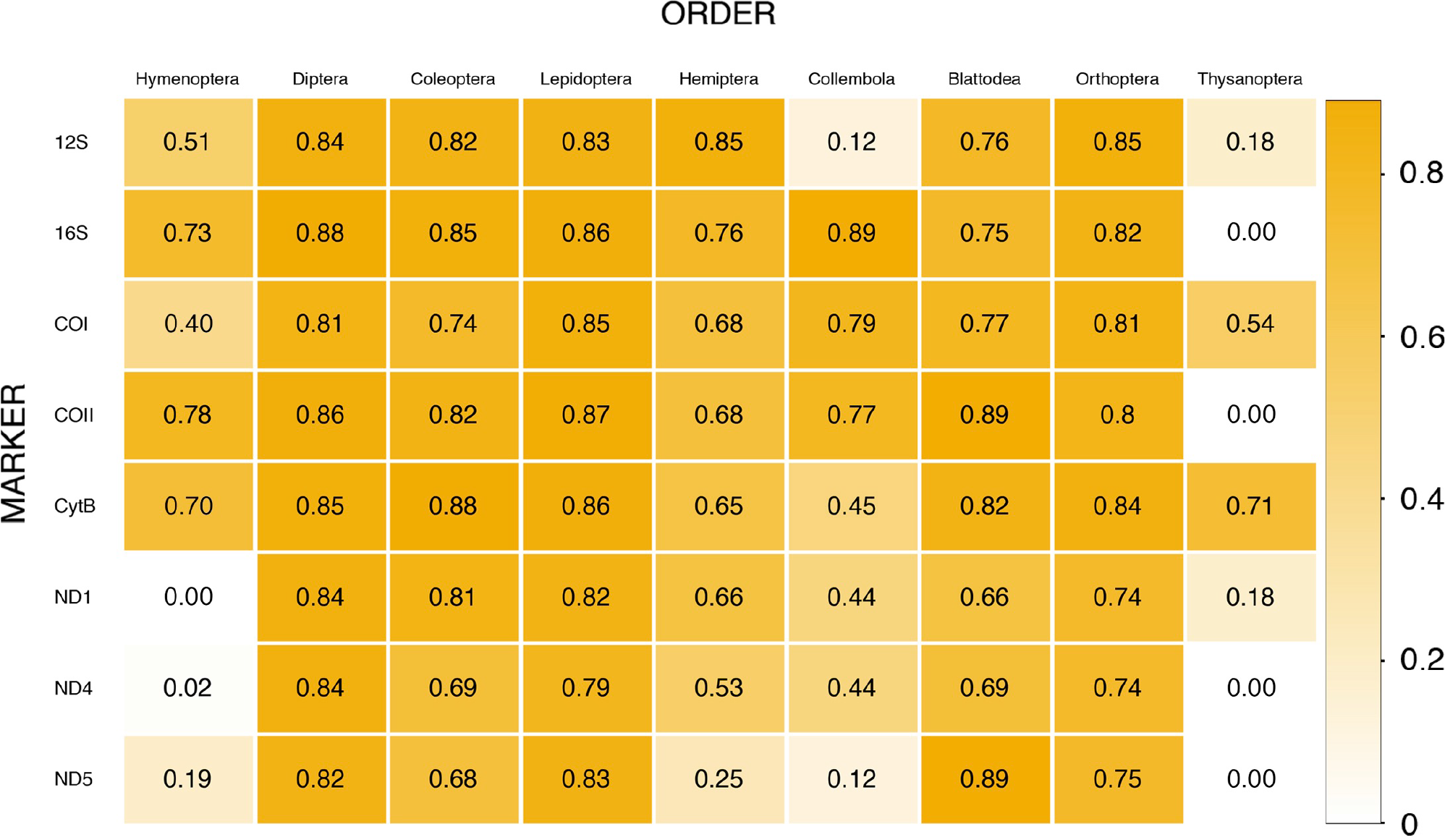
Effective taxonomic resolution of the selected primers for the nine most abundant orders from Hexapoda found in Malaise traps and pitfall traps. The intensity of the color reflects the value.

Looking at the metabarcoding success by insect order instead, the orders that are easiest to analyze (as judged by the average *ETR* across the seven genes) are Diptera, Coleoptera and Lepidoptera, followed by Blattodea, Orthoptera, and Hemiptera, and then by Collembola, Hymenoptera and Thysanoptera (Fig. 4).

### Performance of marker pairs

When combining two markers, the best results were obtained with COI+COII (*ETR* = 0.89), and 12S+COI or 16S+COI (both combinations with *ETR* = 0.88), followed by 12S+16S, 12S+COII, 12S+CytB and COI+CytB (ETR = 0.85) and 16S+COII, 16S+CytB and COII+CytB (*ETR* = 0.84) (Fig. 5A, Tables S10–S11). For the three best marker pairs, which all involve COI, and for COI+CytB, the ETRu of COII, 12S, 16S and CytB is at least double that of COI. Outside these cases, the *ETR*_*U*_ values of the two markers of the pair are more balanced.

**Fig. 5.**
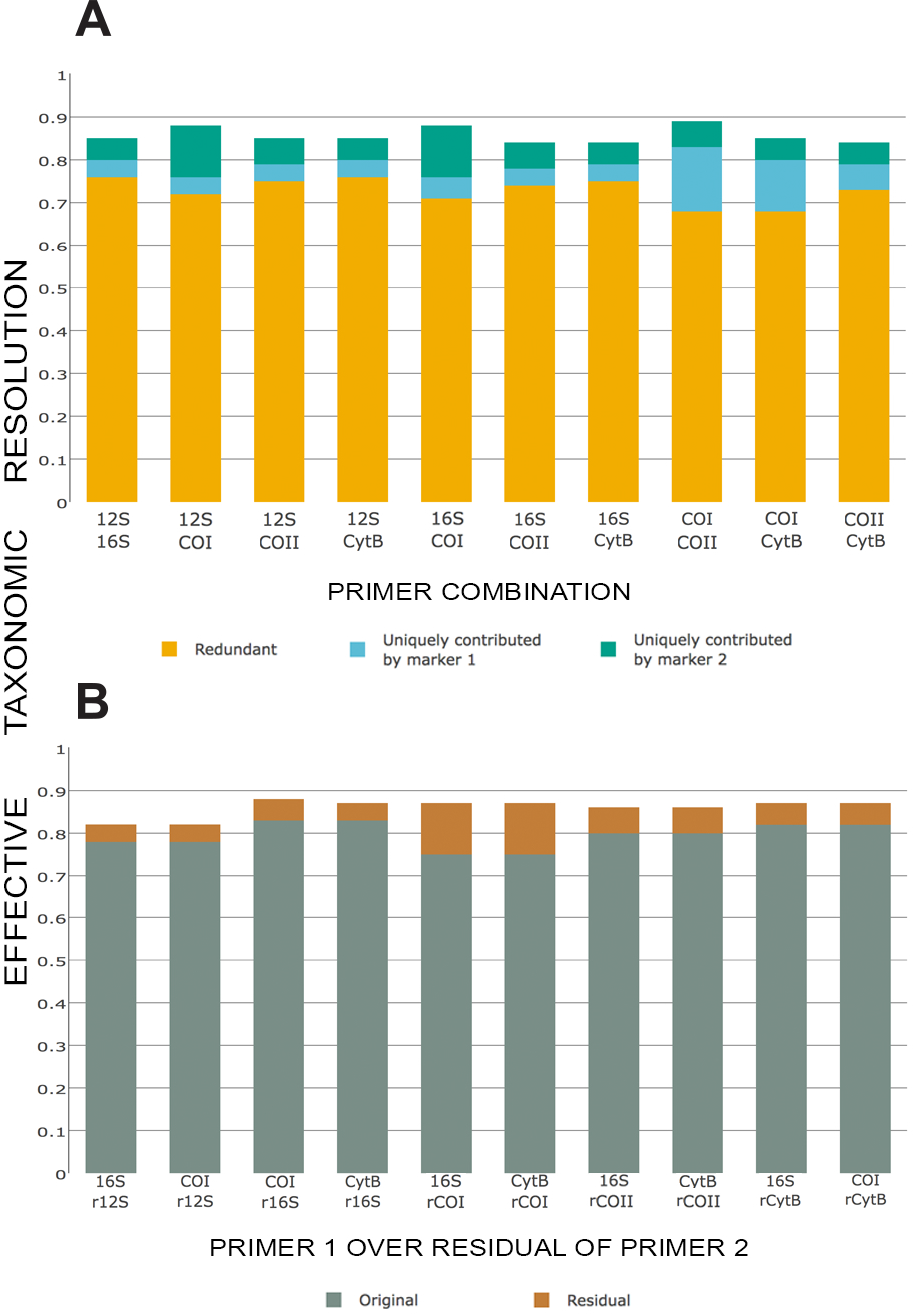
Combined effective taxonomic resolution for all the markers with an *ETR* ≥ 0.75. **A**: simultaneously combined *ETR* showing redundant *ETR* (orange), uniquely contributed *ETR* by the first primer pair (blue) and uniquely contributed *ETR* by the second primer pair (green). **B**: Primer 2 over residual of primer 1; only the best two combinations per original marker are shown (all the combinations are in Fig. S3).

Optimizing the primers for the second marker by designing them only over the set of sequences missed by the first marker (Figs. 5B, S3; Table S9) produced similar results to those obtained by combining the original, independently designed markers (Fig. 5A). The highest *ETR*_*T*_ was obtained when designing a COI primer over the residual of 16S (*ETR*_*T*_ = 0.88), followed by 12Sr16S, 16SrCOI, CytBrCOI, 16SrCOII and CytBrCOII (*ETR*_*T*_ = 0.87). The primers designed for different markers over the residual of 12S obtained the lowest values of *ETR*_*T*_ (0.79-0.82), while the highest were obtained over the residual of 16S (0.86-0.88). In all cases, designing a new primer for one marker over the residual of the same marker (*e.g.* 16Sr16S) provided the lowest values of *ETR*_*T*_ among the combinations for that original marker. In general, the complementary primers designed over residuals are versions of the original primers but with differences in the wobble bases, and the associated amplicons overlap with the original amplicons even though they may differ in length. Only in two cases, COIr12S and 16SrCOI, the amplicons do not overlap with the independently designed amplicons.

## Discussion

The rapid pace in the publication of mitochondrial genomes, and the improvement of *in silico* tools for primer design and evaluation, open up new possibilities in the quest for optimal metabarcoding protocols. Even though the performance of the primers found using *in silico* approaches must still be validated experimentally, these methods are clearly here to stay. The current activity of mitogenome sequencing is well illustrated by the difference in size between the two datasets used for this study, downloaded only eleven months apart and indicating an annual increase of the publicly available mitogenomes by approximately 50 %. The dataset used by us to design primers (D1) was downloaded in September 2015 and contains mitochondrial genomes of more than 800 species. In a previous study published only a year earlier, Clarke *et al.* (2014) had access to data from only 315 insect species (excluding Collembola, Diplura and Protura), less than half of the dataset used here.

One potential problem with the computational approach is that publicly available data do not necessarily reflect the composition of real environmental samples. For instance, Malaise trap samples tend to be dominated by Diptera and Hymenoptera specimens, but these orders are less frequently targeted in mitogenome sequencing projects than more popular groups like Lepidoptera, Coleoptera and some of the minor insect orders. However, the compositional biases are perhaps less problematic than one might fear. Even though they are clearly underrepresented, Diptera and Hymenoptera are still reasonably well represented in the databases we used (Fig. S1).

At a finer taxonomic scale, there may also be important differences between the databases and real environmental samples. For instance, we observed that a few groups, like *Drosophila*, are very well represented in the mitogenome databases, with numerous sequences from the same or very closely related species, while most groups are represented by a diversified selection of mitogenomes from different species. In an environmental sample, such as a Malaise trap sample, one might expect a different type of distribution of sequence abundance and similarity. Nevertheless, given the size of the publicly available databases, we do expect our results to be at least indicative of the performance of the primer pairs in metabarcoding of real samples.

In terms of *in silico* tools, we compared results obtained with ECOPRIMERS, a popular primer design software for metabarcoding studies, with those from DEGEPRIME. Although DEGEPRIME has been widely used for primer design in the field of microbial metabarcoding, its use in eukaryotic studies has so far been restricted to unicellular organisms (Hugerth *et al.* 2014b, Parada *et al.* 2015, Hu *et al.* 2016). However, DEGEPRIME presents a series of advantages over ECOPRIMERS. For instance, it gives the user more control over the parameters that are important in finding adequate primers. It also allows the design of degenerate primers, making it possible to find primers that amplify a larger proportion of the sequences in the database at stringent PCR conditions. In our study, the primers found using ECOPRIMERS did not nearly perform as well as those found with DEGEPRIME, primarily because they were not degenerate.

In the ideal case, a primer pair used for metabarcoding should amplify the desired DNA sequence of all representatives of the target group present in the sample, and bioinformatic processing of these sequences would then be able to identify the species (and their abundance). To do this, the selected DNA sequence should have highly variable regions that are able to discriminate between closely related species, flanked by regions that are conserved across the target group so that they form suitable targets for PCR primers (Ficetola *et al.* 2010). These features should also be present in a short fragment to fit current sequencing platforms, and to allow analysis of degraded DNA (Taberlet *et al.* 2012).

How can these properties be quantified? Ficetola *et al.* (2010; see also Riaz 2011) proposed the indices *B*_*C*_ and *B*_*S*_ for taxonomic coverage and taxonomic resolution, respectively. These indices have proven useful and have been widely employed (Epp *et al.* 2012, Bellemain *et al.* 2010, Clarke *et al.* 2014). However, *B*_*S*_ increases monotonically with the similarity threshold, making it useless in finding the barcoding gap (Fig. 2A; Hebert *et al.* 2003, Meyer & Paulay 2005). Thus, *B*_*S*_ fails to discriminate between inter-and intraspecific genetic variation and does not consider the haplotype diversity within species that barcoding (and, by extension, metabarcoding) should assume. Only the presence of a rich reference database would allow safe identification of the clusters that belong to the same species, and even the COI reference databases are still not complete enough for this in most cases. Therefore, *B*_*S*_ is not an adequate measure for comparing the performance of metabarcoding markers.

In theory, the exclusive taxonomic resolution index we propose here (*B*_*E*_) solves these problems. Given a rich reference database covering intraspecific variation in all taxa, it should allow us both to identify the barcoding gap and to compare the performance of different metabarcoding markers when species circumscriptions cannot be deduced from a reference database. However, we found that *B*_*E*_ did not decrease rapidly enough at high similarity values to allow safe identification of the barcoding gap using our database. The reason is apparently the small number of species for which any intraspecific variation is covered in our database. Therefore, we also propose the alternative definition of the index, 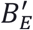, which measures resolution relative to clusters and not to species in the database. This results in the index being increasingly penalized as the few abundantly represented species are split into smaller and smaller clusters when the similarity threshold value is increased beyond the barcoding gap. This version of the index allowed us to easily find the barcoding gap (Fig. 2). Our modified index should lag slightly behind *B*_*E*_ in the decrease of resolution seen beyond the barcoding gap because of the relative shortage of high amounts of intraspecific diversity in the database. However, the decrease in 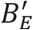 should be faster and more dramatic at high threshold values than for *B*_*E*_, as is also evidenced by our plot (Fig. 2). At the barcoding gap, 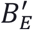 should be a good approximation of *B*_*E*_.

We asked ourselves why we never obtained exclusive taxonomic resolution over 0.90 in our analyses, even for COI markers that supposedly should provide almost perfect taxonomic resolution. The fact that most (67-86 %) of the unresolved species were shared between marker pairs (Table S7) suggests that most of these species are impossible to circumscribe correctly using the mitochondrial data in the database. This could be because the taxonomic annotation in the database is incorrect, or because the mitochondrial genetic variation within species is unusually small or high in these species. In either case, it seems likely that the taxonomic resolution values presented here are conservative estimates of the true values.

Arguably, the best metabarcoding primers we found are the 12-fold degenerate primer pairs targeting 16S and 12S. It was necessary to increase the degeneracy level considerably to obtain similar performance for the primers targeting protein-coding genes. Clearly, this reflects the superiority of rRNA genes as metabarcoding markers because of the presence of continuous, highly conserved regions allowing the design of universal primers. The variability seen at third codon positions in protein-coding genes makes it more difficult to find good primers. There is also an increased risk that new sequence variants not considered in the design phase will show up in environmental samples, and that they will not be amplified due to primer mismatch. One might have expected the performance of rRNA markers to be negatively affected by the difficulty of separating closely related species based on a conservative sequence, but our results indicate that this is a not a major issue. The exclusive taxonomic resolution of the best rRNA markers is very similar to that of the best protein-coding markers. Closely related species have more similar rRNA marker sequences, but they are still distinct.

Boosting the performance of metabarcoding markers in protein-coding genes using highly degenerate primers can potentially present several problems, such as higher risk of primer dimers or of binding of primers to off-target regions in the genome. Nevertheless, we focused our detailed studies of protein-coding markers on 216-fold degenerate primers because of their vastly superior performance *in silico* compared to 12-fold degenerate primers. Primers targeting conservative rRNA genes also come with a risk, namely that they will amplify a large number of sequences of nontarget taxa, such as bacteria that may be present in water or soil samples used in arthropod inventories. For these reasons, we chose to focus our analyses on rRNA primers with 12-fold degeneracy, even though there was a modest but still significant increase in the ETR values of the rRNA primers with 216-fold degeneracy. These choices should be borne in mind when interpreting our results.

Although our results show that several other protein-coding genes (notably COII and CytB) offer excellent markers for metabarcoding at 216-fold degeneracy, the COI results are of particular interest because of the rich reference data available for this gene. The best primer pair we found for COI was the one amplifying the BE fragment, with the primers ArF2 - ArR5 (Gibson *et al.* 2014), with an ETR of 0.84, followed by the pairs BF2 - BR2 (Elbrecht & Leese 2017) (*ETR* = 0.78), HexCOIF4 - HexCOIR4 (*ETR* = 0.75) and BF2 - BR1 (Elbrecht & Leese 2017) (*ETR* = 0.75). However, it must be noted that the high *ETR* value for the BE fragment is only obtained when inosine is considered to pair with all other bases. Inosine actually only binds to adenine, thymidine and cytosine; if this is taken into account, the *ETR* of this marker decreases to 0.65, lower than the other primer pairs. The next best primer pair is BF2 - BR2, but the associated amplicon (422 bp) is longer than is ideal for current HTS platforms. Thus, the best COI primers that we found are thus the pairs BF2 - BR1 and HexCOIF4 - HexCOIR4. These two pairs are almost identical, with only two different bases in the reverse primer, which at least *in silico* grant HexCOIF4 - HexCOIR4 a slightly higher *ETR* value. The first of these primer pairs was presented only very recently, the second is new to this study. The primers used in most of the published COI metabarcoding studies of insect diversity have considerably lower performance than these primers according to our results.

Our taxon-specific analysis confirms previous studies (Clarke *et al.* 2014, Elbrecht *et al.* 2016), which have shown that 16S markers present considerably less amplification bias than COI markers across insect taxa. Nevertheless, 16S markers fail to detect Thysanoptera. Markers in the 12S, COI, COII and CytB genes are mostly able to detect Thysanoptera but they show considerably more variation in ETR across other insect orders. Due to its high degeneracy, the best COI primer pair studied here overcomes most of the bias reported previously for other COI metabarcoding primers (Yu *et al.* 2012, Clarke *et al.* 2014, Brandon-Mong *et al.* 2015). In conclusion, no marker has perfect coverage across all insect orders, and a combination of primers should be considered if broad taxonomic coverage is needed.

Our results indicate that it is possible to increase the effective taxonomic resolution from about 0.80 to almost 0.90 by combining two markers. As noted above, this is close to the maximum resolution obtainable for the database. The best combinations are arguably 12S-COI, 16S-COI and COI-COII, since they show high *ETR*_*T*_ values while allowing one to take advantage of the extensive reference data for COI. The combinations 16S-COI and COI-COII are especially interesting for insect metabarcoding with broad taxonomic coverage, since both 16S and COII provide high *ETR* for Hymenoptera (the most diverse order of Hexapoda). Designing the second primer pair over the sequences not amplified by the first primer pair did not significantly improve the results. Presumably, this occurs because the residual sequences still represent a large portion of the original taxonomic and genetic diversity. Desigining complementary primers over residuals may nevertheless be useful for well-known biotas, especially for targeting the specific taxa that COI cannot detect. However, for poorly documented areas, there is a significant risk of missing taxa that are not amplified by the original marker but are not present in the primer design database either. In such cases, it may be better to use independently designed primers.

Multi-locus or multi-gene approach has been used in several metabarcoding studies as a solution to fill the gaps in the detection of species by a single marker that is not able to amplify or discriminate between certain species (Cowart *et al.* 2015, Drummond *et al.* 2015, Shaw *et al.* 2016). Alberdi *et al.* (2017) pointed out that the use of different markers targeting the same taxon reduces the effect of the biases of individual primer sets and increases the taxonomic coverage of the sample. In insect surveys, however, the multi-locus approach has been less frequently applied (Elbrecht *et al.* 2016, Alberdi *et al.* 2017).

Even though the use of two primer pairs can increase taxonomic resolution significantly, it must be borne in mind that the individual HTS reads of the two markers cannot be combined, at least not using standard protocols. Thus, advanced bioinformatic processing will be needed to synthesize results across analyses using two separate markers. Interestingly, such analyses could help improve the quantification of the abundance of different species if amplification biases differ between markers.

A key finding of this study is that the best metabarcoding markers for insects are not found in the COI gene. The rRNA markers offer much broader taxonomic coverage at low levels of primer degeneracy and under stringent PCR conditions, while still resolving most species that can be separated genetically. At high levels of primer degeneracy, markers in the protein-coding genes can compete in performance with the rRNA markers, but under such conditions the best COI markers are often outperformed by markers in other genes like COII and CytB. It is true that we currently lack reference data for these alternative markers. However, there is much to suggest that recent advances in the sequencing of mitochondria, such as mitochondrial metagenomics (Zhou *et al.* 2013, Crampton-Platt *et al.* 2016, Ciccionardi *et al.* 2017) and long-range PCR for whole mitochondrial genome amplification (Deiner *et al.* 2017), will change this in the near future. As the public mitogenome databases grow in size, so do the reference libraries for all mitochondrial markers, as the existing COI reference data can be used to provide reliable taxonomic annotation of entire mitochondrial genomes, where such annotation is not available from other sources. HTS techniques can also now be used to facilitate the generation of reference libraries for custom markers during local or national barcoding campaigns. Therefore, we think metabarcoding projects should seriously consider the use of markers outside of COI as a complement to or replacement of COI-based analyses.

## Acknowledgements

We are thankful to Niclas Gyllenstrand for asking the question that resulted in the formalization of the combined ETR indices, and to Chiara Leo for her assistance in the curation of the mitochondrial datasets. This project was funded by the European Union’s Horizon 2020 research and innovation programme under the Marie Skłodowska-Curie grant agreement No. 642241 (BIG4 project, http://big4-project.eu) and by a grant from the Swedish Research Council (No. 2014-05901).

## Author contributions

DM, AFA and FR conceived and designed the study. DM collected the data, conducted the analysis, wrote the first draft of the paper, and prepared the figures and tables. DM, AFA and FR reviewed and contributed to subsequent drafts of the manuscript.

## Supplementary material

**Table S1** Taxonomic composition of the first set of mitochondrial genomes (downloaded October 2015) (.csv file).

**Table S2** Taxonomic composition of the second set of mitochondrial genomes (downloaded September 2016) (.csv file).

### Supplementary material: Tables

**Table S12** Primers designed using DEGEPRIME, max degeneracy = 12-fold (.csv file).

**Table S13** Primers designed using DEGEPRIME, max degeneracy = 216-fold (.csv file).

**Table S14** Primers designed using ECOPRIMERS (.csv file).

**Table S15** Previously published primers (.csv file).

**Table S16** Primers for residual species designed with DEGEPRIME (.csv file).

**Supplementary material: Figures**.

